# Resolution of MALDI-TOF MS Compared to Whole Genome Sequencing for the Identification of *Vibrio parahaemolyticus* Strains Isolated from Oysters

**DOI:** 10.64898/2026.06.11.731630

**Authors:** Tetyana King, Mayela Pedrueza, Maliha Rahman, Bonnie Oh, Michael G. LaMontagne

## Abstract

Climate change and eutrophication are driving the expansion of the range of *Vibrio* species, including *V. parahaemolyticus*. This bacterium is a major foodborne pathogen and understanding the biogeography of virulent strains of this species is crucial for ensuring food safety. Whole-genome sequencing (WGS) provides strain-level identification of bacteria and is widely used for tracking bacterial pathogens; however, WGS is costly and labor-intensive. Matrix-assisted laser desorption/ionization time-of-flight mass spectrometry (MALDI-TOF MS) provides a rapid, accurate, and cost-effective method for bacterial identification; however, the resolving power of MALDI-TOF MS and WGS for *V. parahaemolyticus* has not been systematically compared. In this study, 70 *V. parahaemolyticus* strains were isolated from oysters (*Crassostrea virginica*) collected from the Gulf Coast and Massachusetts. Oysters were collected in Galveston Bay (Texas) and aquaculture plots in Massachusetts, and purchased from seafood markets in Texas and Louisiana in the U.S. For comparison, two isolates of *V. anguillarum* were cultured from the exoskeleton of blue crabs purchased from a seafood market in Seabrook (Texas). All isolates were identified using the MALDI Biotyper system and analyzed with custom R scripts. Cluster analysis of mass spectra generated by MALDI-TOF MS, and phylogenomic analysis revealed distinct clusters corresponding to the source of oysters. In both the mass spectra and WGS analysis, *V. parahaemolyticus* strains isolated from Massachusetts formed a coherent cluster. For comparisons between species, cosine similarities of mass spectra generated by MALDI-TOF MS ranged from 0.43 to 0.59, and average nucleotide identity (ANI) values generated by WGS ranged from 76% to 77%. For comparisons within species, cosine similarities of mass spectra ranged from 0.68 to 0.91 and ANI values ranged from 98% to 100%. This suggests that MALDI-TOF MS has a resolution comparable to WGS and can be used to track strains of *V. parahaemolyticus* associated with oysters.

## 1. Introduction

Climate change and the global trophic state change (distribution of dissolved organic carbon, ammonium, chlorophyll a, silicate and carbon/nitrogen ratio of suspended organic particulates) has expanded the habitat of bacteria from the *Vibrionaceae* family, including pathogenic and antibiotic-resistant strains (Kopprio et al. 2017; Semenza et al. 2017; Vezzulli et al. 2016). This expansion of range has doubled the size of the human population at risk for *Vibrio* infections (Trinanes & Martinez-Urtaza 2021), including the risk from consuming seafood (Ndraha & Hsiao 2022).

*V. parahaemolyticus* is commonly found in coastal waters where temperatures exceed 18°C and salinity is low (<25 ppt NaCl) (Baker-Austin et al. 2013). This Gram-negative bacterium is among the most common foodborne pathogens (Alipour et al. 2014; Letchumanan et al. 2014) and highly virulent strains of *V. parahaemolyticus* have recently emerged (Lee et al. 2015). Similar to many *Vibrio* species, *V. parahaemolyticus* can adhere to the surface of marine animals and form a biofilm, which enhances bacterial survival (Gode-Potratz & McCarter 2011) and causes significant damage to the aquaculture industry (Letchumanan et al. 2014; Letchumanan et al. 2015). Infections of humans typically occur through the consumption of undercooked or raw seafood, including shellfish, fish, and crabs (Daniels et al. 2000; McLaughlin Joseph et al. 2005). Rapid and accurate identification of *V. parahaemolyticus* is essential for epidemiological surveillance and reducing public health risks (Canellas et al. 2024). Identification of *V. parahaemolyticus* in oysters is crucial. These bivalves are filter feeders that concentrate bacteria from the environment, and since oysters are often consumed raw or undercooked, they represent a direct path for transmission of pathogenic bacteria to humans (Banerjee et al. 2025).

Identification of virulent strains of bacteria is necessary to track outbreaks and biogeographical patterns. Whole genome sequencing (WGS) provides high-resolution, comprehensive genetic data that enables accurate species identification and strain level differentiation. In addition, WGS is used for the detection of virulence factors and antimicrobial resistance (AMR) genes, offering critical insight into pathogenic potential of bacteria and the risk of dissemination of foodborne pathogens (Gomes et al. 2025). The high discriminatory power of WGS supports bacteria source tracking that is important for epidemiological investigations (Brown et al. 2021) and can identify clusters of *V. parahaemolyticus* strains isolated from oysters harvested from different locations (Miller et al. 2021) However, WGS is expensive and time-consuming (Price et al. 2023). In contrast, matrix-assisted laser desorption/ionization time-of-flight mass spectrometry (MALDI-TOF MS) is a fast, accurate, and relatively inexpensive method for bacterial identification (Clark et al. 2013; Singhal et al. 2015).

The accuracy of MALDI-TOF MS has been validated for a wide range of Gram-negative bacteria, including *Cronobacter* spp. (Stephan et al. 2010), *Shigella* spp. (Tan et al. 2012), *Enterobacter* spp. (Pavlovic et al. 2012), *Aeromonas* spp. (Donohue et al. 2006), *Vibrio* spp (Boonstra et al. 2023; Vidal et al. 2020) and others (Couderc et al. 2012; de Jong et al. 2013). Several studies have included *V. parahaemolyticus* (Banerjee et al. 2025; Hazen et al. 2009; Li et al. 2018; Sulaiman et al. 2023). Cluster analysis of mass spectra generated by MALDI-TOF established that MALDI-TOF MS has a higher resolution than 16S rRNA gene sequencing for differentiating *V. parahaemolyticus* from closely related species (Mougin et al. 2020) and MALDI-TOF MS can reliably identify *V. parahaemolyticus* and differentiate this species from other *Vibrio* species isolated from seafood (Banerjee et al. 2025), but MALDI-TOF MS can show incoherent clusters for this species (Canellas et al. 2024). To our knowledge, the comparative resolving power of MALDI-TOF MS and WGS has not been evaluated for *V. parahaemolyticus*strains isolated from oysters.

In this study, we compared the resolution of MALDI-TOF MS and WGS for tracking the source of *V. parahaemolyticus* in oysters. Libraries of isolates were generated from oysters collected from aquaculture facilities, harvested wild, and purchased from seafood markets. Draft genomes of 70 isolates were assembled using Illumina sequencing, and phylogenetic relationships were inferred with a maximum likelihood tree based on alignment of core genomes. For comparison, complete genomes and mass spectra were also generated from two isolates of *V. anguillarum*. The resolution of MALDI-TOF MS was expressed as the cosine similarity coefficient and calculated using custom R scripts. Average nucleotide identity (ANI) was determined based on WGS, which served as the reference method to estimate the accuracy of inter-species and within-species identification of *V. parahaemolyticus*. MALDI-TOF MS and WGS gave comparable results, in terms of identifying coherent clusters of *V. parahaemolyticus* isolates that corresponded to the source of the oysters. This suggests that MALDI-TOF MS can distinguish strains within the species *V. parahaemolyticus*.

## 2. Materials and Methods

### 2.1 Bacteria Collection and Culturing

*V. parahaemolyticus* isolates were collected from eastern oysters (*Crassostrea virginica*) harvested in Galveston Bay (Texas) (August 2024 and October 2024), aquaculture plots in Massachusetts (July 2024), and purchased from seafood markets in Louisiana (March 2024) and Texas (April 2023). Oysters from Texas and Louisiana were transported on wet ice immediately to the University of Houston – Clear Lake and processed within six hours of collection. Oysters from Massachusetts were shipped overnight on ice packs and processed within 48 hours of collection. Samples were prepared following the Bacteriological Analytical Manual of the FDA (USA) (Kaysner et al. 2004). Briefly, oyster tissue was mixed with 100 mL of alkaline peptone water (APW) (peptone powdered, cat. # 20-260, APEX, El Cajon, CA, USA) and homogenized for 3 min in a Waring blender. The resultant slurry was serially diluted to 10^-3^ in APW and 200 µl of the dilutions were dispensed in sterile 96-well plates (32 wells for 10^-1^, 10^-2^, and 10^-3^) and incubated at 30°C for 24 h. Bacteria were transferred from positive wells to thiosulfate-citrate-bile-sucrose agar (TCBS, cat. # M189, HiMedia, Maharashtra, India) plates at 30 °C for 24 h. Isolated colonies were then restreaked on marine agar (MA) (M384, HiMedia) and incubated at 30 °C for 24 h.

*V. anguillarum* isolates were collected from the exoskeleton of a blue crab (*Callinectes sapidus*) purchased from a seafood market in Texas in February 2023 following the procedure described previously (Davis & Sizemore 1982) with the following modifications. Briefly, the ventral surface of the lateral spine of the crab was sampled with a sterile swab. The swab was placed into a 1.5 mL centrifuge tube with 1000 µL phosphate-buffered saline solution (PBS, pH 7.5, cat. # E703, VWR, Radnor, PA, USA) and thoroughly vortexed. This solution was diluted to 10^-3^ in PBS and 10 µL was transferred into MA plates and incubated 30 °C for 24 h. Isolated colonies were selected, re-streaked onto secondary MA plates, and incubated at 30 °C for 24 h before MALDI-TOF MS analysis.

### 2.2 MALDI-TOF MS

Isolates were prepared for MALDI-TOF MS using the full extraction method recommended by Bruker Scientific (Billerica, MA, USA). Briefly, ethanol deactivation of bacteria was performed by transferring fresh single colonies to 1.5 mL centrifuge tubes with 300 µL LC–MS water (OmniSolv™, cat. # WX001, EMD Millipore Corporation, Burlington, MA, USA), vortexing, and adding 900 µL ethanol (OmniPur Ethyl Alcohol, cat. # 4450, Sigma, St. Louis, MO, USA). The tubes were vortexed and centrifuged at 13,000 x g for 2 min. After pouring out the supernatant, the tubes were centrifuged at 13,000 x g for 1 min and residual ethanol was removed. The pellet was dried in open-lid tubes at room temperature (RT) for 15 min before it was dissolved in 50 μl of 70% formic acid (Cat. # 33015, puriss Sigma) and incubated at RT for 5 min, then 50 μl of acetonitrile (Cat. # 900667, Sigma) was added. The tubes were thoroughly vortexed, incubated at RT for 5 min and centrifuged at 13,000 x g for 2 min. The supernatant was transferred to clean 1.5 ml tubes and stored at -20°C for less than 2 weeks before MALDI-TOF MS analysis.

### 2.3 Mass spectra acquisition

For MALDI-TOF analysis, 1 μl of extracted protein solution was spotted on a polished steel target (Cat. # 8280800, Bruker Scientific). Two technical replicates of each isolate were used for identification of bacteria. Eight technical replicates were spotted for main spectra profile creation, and each spot was measured three times to generate 24 spectra per isolate. Bacterial test standard (BTS) (Cat. # 8255343, Bruker Scientific) was dissolved in 50 μl Bruker standard solvent (Cat. # 900666, Sigma) and placed on two spots. The target was dried at RT for 10 min and then each spot was overlaid by 1 μl of fresh matrix solution, which was prepared by dissolving 0.01 g of 10 mg/mL a-cyano-4-hydroxycinnamic acid (Cat. # 476870, Sigma) in 100 μl of Bruker standard solvent. The target was dried at RT for matrix crystallization and immediately used for spectra acquisition.

Mass spectra were acquired using a positive polarity MALDI Biotyper mass spectrometer (Biotyper Sirius v1.6.0.880 Bruker). Mass spectra were obtained under MBT_autoX method with the mass range from 2000 to 21000 Da, linear detector gain (31x 2533 V) with a laser frequency of 200 Hz and 390 ns of pulse ionization. Ion sources were 19.84 kV for the first and 18.13 kV for the second objective was 5.96 kV. Smartbeam parameter was in default mode. Real-time smoothing was disabled, baseline shift adjustment was set to 0% and analog offset was set to -0.7 mV. Each spot was measured with a resolution of 0.5 G/s. Before spectra acquisition, the system was automatically calibrated using BTS calibration proteins in the mass range from 3637.8 Da to 16952.3 Da. Peak assignment tolerance was 1000 ppm. The identification of isolates was performed using flexControl software (v3.4) (Bruker) and the default MALDI Biotyper library (v11.0.0.0) (Bruker), containing main spectra profiles (MSP) of 54 *Vibrio* spp, and *V. parahaemolyticus* was represented by nine MSPs (Liu et al. 2022). A log score of 2.00 to 3.00 correlated with high identification confidence at the species level, and 1.70 to 1.99 at the genus level (File S1).

### 2.4 Custom main spectra profiles creation

MSPs of *V. parahaemolyticus* were created following Bruker recommendations. Briefly, 24 spectra per isolate were manually evaluated with the FlexAnalysis Software (v3.4) using the MBTstandard FAMS Method. Spectra were smoothed, and baseline was subtracted. Flat spectra and outliers were removed from the dataset. Spectra that passed quality control were checked for peak shifts (500 ppm). At least 21 spectra per isolate were used for custom MSP creation with the automatic function in MALDI Biotyper Compass Explorer (Bruker, v4.1.100).

### 2.5 Whole genome sequencing

#### V. parahaemolyticus

At the Texas Public Health Laboratory, *V. parahaemolyticus* cultures were streaked for purity on Trypticase Soy Agar (5% sheep’s blood) plates and incubated at 37℃ for 18-24 hours overnight. DNA was extracted using the MagNA PURE 96 DNA and Viral NA Small Volume Kit (Cat. # 06543588001, Roche) on the MagNA PURE 96 automated instrument. Libraries were prepared with the Illumina DNA Prep Kit (Cat. # 20060059, Illumina) and sequenced on an Illumina MiSeq platform using v3 reagents and Illumina NextSeq 2000 platform using XLEAP-SBS P1 reagents.

The genomes for *V. parahaemolyticus* isolates were de-novo assembled with SKESA (v2.2) (Souvorov et al. 2018). Quality control of the reads and draft genomes, of *V. parahaemolyticus* isolates was conducted with the GalaxyTrackr MicroRunQC workflow, as described previously (Timme et al. 2025).

#### V. anguillarum

Genomic DNA from the two *V. anguillarum* isolates was extracted using MasterPure Complete DNA and RNA Purification Kit (Cat. # MC85200, LCG Biosearch Technologies, Teddington, UK) for DNA extraction following the manufacturer’s instructions. DNA was purified using the GenElute-E Single Spin DNA Cleanup Kit (Cat. # EC600, Sigma). The library was prepared with the Nextera DNA Flex Library Prep Kit (Cat . # 20015829; Illumina) and sequenced on an Illumina MiSeq platform using v3 reagents. Long-read sequencing was performed using the MinION Mk1C (Oxford Nanopore Technologies (ONT), Oxford, UK) with the Native Barcoding Sequencing Kit (Cat. # SQK-NBK114.24, version NBE_9196_v114_revQ_15Sep2022), following the manufacturer’s instructions. The library was sequenced for 48 hours using FLO-MIN106 R9 flow cell connected to MinKNOW software (v24.02.08; ONT).

Raw read quality was assessed using Fastp (v0.23.2) (Chen 2023) and trimmed with Trimmomatic (v0.39; https://github.com/usadellab/Trimmomatic). The hybrid genome sequence assembly was performed with Unicycler pipeline (v0.5.0) (Wick et al. 2017), and CheckM (v1.2.2) was used to check the quality of WGS (Parks et al. 2015).

### 2.6 Data analysis

Phyloproteomic analysis of MALDI-TOF mass spectra was conducted using a custom script (Mazhari et al. 2025) written in R (v4.4.1). The script incorporated functions from MALDIquant (v1.22.3) and MALDIquantForeign (v0.14.1) packages (Gibb & Strimmer 2012), along with PVclust (v2.2.0) for hierarchical clustering with the approximately unbiased p-value and bootstrap probability support (Suzuki & Shimodaira 2006). Spectra were processed and analyzed with two loops. The preliminary analysis loop was run with 200 iterations for selection optimal values for processing parameters, including smoothing with Savistky-Golay filter, baseline subtraction with SNIP method, and signal-to-noise ratio (SNR). Low-quality spectra were removed. Internal calibration of spectrum was performed using reference peaks in m/z range of 2833-9552 Da. Pairwise similarity was obtained from cosine and Jaccard coefficients calculated using Philentropy (v0.9.0) package (Drost 2018). The second loop, with 1000 iterations, used the optimize parameters for testing MALDI-TOF Taxonomic Units (MTUs) coherence (LaMontagne et al. 2021) using RWeka (v0.4-46) (Hornik et al. 2009) and performed hierarchical clustering with PVclust.

ANI between genomes of *V. parahaemolyticus* and *V. anguillarum* and reference genomes for each species, and between pairs of isolates were calculated with FastANI (v1.33) (Jain et al. 2018). Prokka (v1.14.6) (Seemann 2014) was used for genome annotation, and Roary (v3.13.0) (Page et al. 2015) was used for pan-genome analysis and core core-genome alignment with a strict core-gene definition (-e) and without paralog splitting (-s). Core genes were aligned using MAFFT (--mafft) (v.7.490). IQ-Tree (v2.4.0) was applied to the resulting concatenated core gene alignment (Nguyen et al. 2014). The gene presence/absence matrix created with Roary was used for rarefaction analysis of clusters of orthologous groups (COG). Using a custom R script, random sampling was performed 10 times for each number of isolates and unique COGs were calculated. The maximum-likelihood phylogenetic tree was created based on core gene analysis. Additional genomes collected from oysters in Florida (R17FL, R126FL, R130 FL), Louisiana (R13LA, R33LA), Maine (R59ME), South Caroline (R135SC), Texas (R21TX), and Virginia (R74VA) and originally published by Miller and colleagues (Miller et al. 2021) are incorporated into the tree for increasing phylogenetic resolution. Genomes of *V. anguillarum* (CS13 and CS14) were not included in the analysis. The tree was visualized using iTol Online (v7) (Letunic & Bork 2021).

*V. anguillarum* genomes (CS13 and CS14) were annotated and visualized using Bakta (v1.11.0) (Schwengers et al. 2021). The number of protein-coding sequences (CDS) was predicted with Prodigal (v2.6.3) (Hyatt et al. 2010) and GeneMarkS (v4.28) (Besemer et al. 2001). tRNAscan-SE (v2.0.9) (Chan & Lowe 2019) and Barrnap (v0.9; https://github.com/tseemann/barrnap) were used to predict numbers and positions of tRNA and rRNA sequences. Tandem repeats were found and annotated with Tandem Repeats Finder (v4.09) (Benson 1999), RepeatMasker (v4.1.9) (Tempel 2012), and RepeatModeler (v2.0.7) (Flynn et al. 2020).

Genes associated with antimicrobial resistance (AMR) and virulence (VF) for all genomes were predicted with resistance gene identifier (RGI) (v6.0.4) using CARD database (v3.2.7) (Alcock et al. 2023) and abricate (v1.0.1) with Virulence Factor Database (VFDB) (https://github.com/tseemann/abricate).

## 3. Results

All draft genomes, generated from the 70 *V. parahaemolyticus* isolates studied herein, showed high similarity, in terms of ANI, to the reference genome for *V. parahaemolyticus* RIMD1210633 (RefSeq ID: GCF_000196095.1). ANI values between the reference genome and the 70 isolates included in this study exceeded 98% for all comparisons (File S1). All 70 of these isolates were identified as *V. parahaemolyticus* by matching mass spectra, generated by MALDI-TOF MS, to the database in Bruker system. All of these matches had scores > 2.3, which is the cutoff recommended by the manufacturer for highly confident identification at the species-level (File S1).

The draft genomes assembled de novo for the 70 isolates of *V. parahaemolyticus* ranged from 5.0 to 5.9 Mb in size, with an average GC content of 45% (File S1). The number of coding protein genes ranged between 4,609 and 5,444, which corresponds to previously described genome assemblies for this species (Bosi et al. 2024; Liu et al. 2024; Prithvisagar et al. 2021). The largest genome - 5,916,921 bp - was estimated for isolate KM44 (seafood, Texas), the smallest genome - 5,022,921 bp - was estimated for isolate YK510 (wild oyster, Texas) (File S1). Pan-genome analysis of 70 *V. parahaemolyticus* isolates identified 14,867 genes, of which 2,686 genes, or approximately 18% of the total, were assigned to core genes and were present in ≥ 99% of studied isolates. AMR profiles included β-lactam, trimethoprim, and tetracycline resistance genes. Virulence-associated genes were grouped into functional categories, including regulators and accessory factors, as well as components of the Type III secretion system (T3SS). These clusters were primarily involved in host cell interaction and pathogenicity, with T3SS mediating the delivery of effector proteins into host cells, while accessory factors and regulators contribute to virulence regulation and environmental adaptation (Galán & Collmer 1999; Makino et al. 2003). Virulence-associated clusters were largely conserved across *V. parahaemolyticus* isolates, suggesting similar virulence potential (File S2).

Species-level identification of *V. anguillarum* isolates CS13 and CS14 (File S3) was confirmed by ANI analysis against the reference genome *V. anguillarum* PF4-E1-1 (RefSeq ID: GCF_003390515.1). The closest related genome, as assessed by a BLAST of the NCBI database, was *V. anguillarum* strain MHK3 (accession number CP022468), which was isolated from a flounder in China. The AMR profile of CS13 and CS14 included vancomycin resistance genes. Virulence-associated genes were grouped into functional categories, including regulators, motility and chemotaxis systems, toxin and virulence-associated systems, and iron acquisition systems. These clusters were largely conserved between isolates CS13 and CS14, consistent with their shared AMR and virulence gene profiles, and were associated with host colonization, environmental adaptation, and pathogenicity (Frans et al. 2011; Romeo et al. 2013) (File S4). Both *V. anguillarum* isolates had two circular chromosomes, as expected (Naka et al. 2011) (Files S5–S8).

Phylogenetic analysis of *V. parahaemolyticus* genomes, performed on the core genome alignment, showed that isolates collected from aquaculture oysters from Massachusetts (MS09–MS25) (File S1) form a tight, distinct monophyletic clade (Fig. 1). Most isolates from wild oysters from Galveston Bay (YK543–YK565 and YR19–YR22) also form coherent clusters. In contrast, isolates from oysters purchased from seafood markets in Texas and Louisiana do not form stable clades and show genetic relatedness to *V. parahaemolyticus* genomes collected from oysters in Florida (R17FL, R126FL, R130 FL), Louisiana (R13LA, R33LA), Maine (R59ME), South Carolina (R135SC), Texas (R21TX), and Virginia (R74VA). These genomes were originally published by Miller et al. (Miller et al. 2021). Notably, isolates KM03 and KM04 (seafood market, Texas), which have distinct MALDI-TOF proteome profiles, were almost identical in genetic content. These two genomes are represented as KM03 in the phylogenetic tree (Fig. 1).

**Figure 1.**
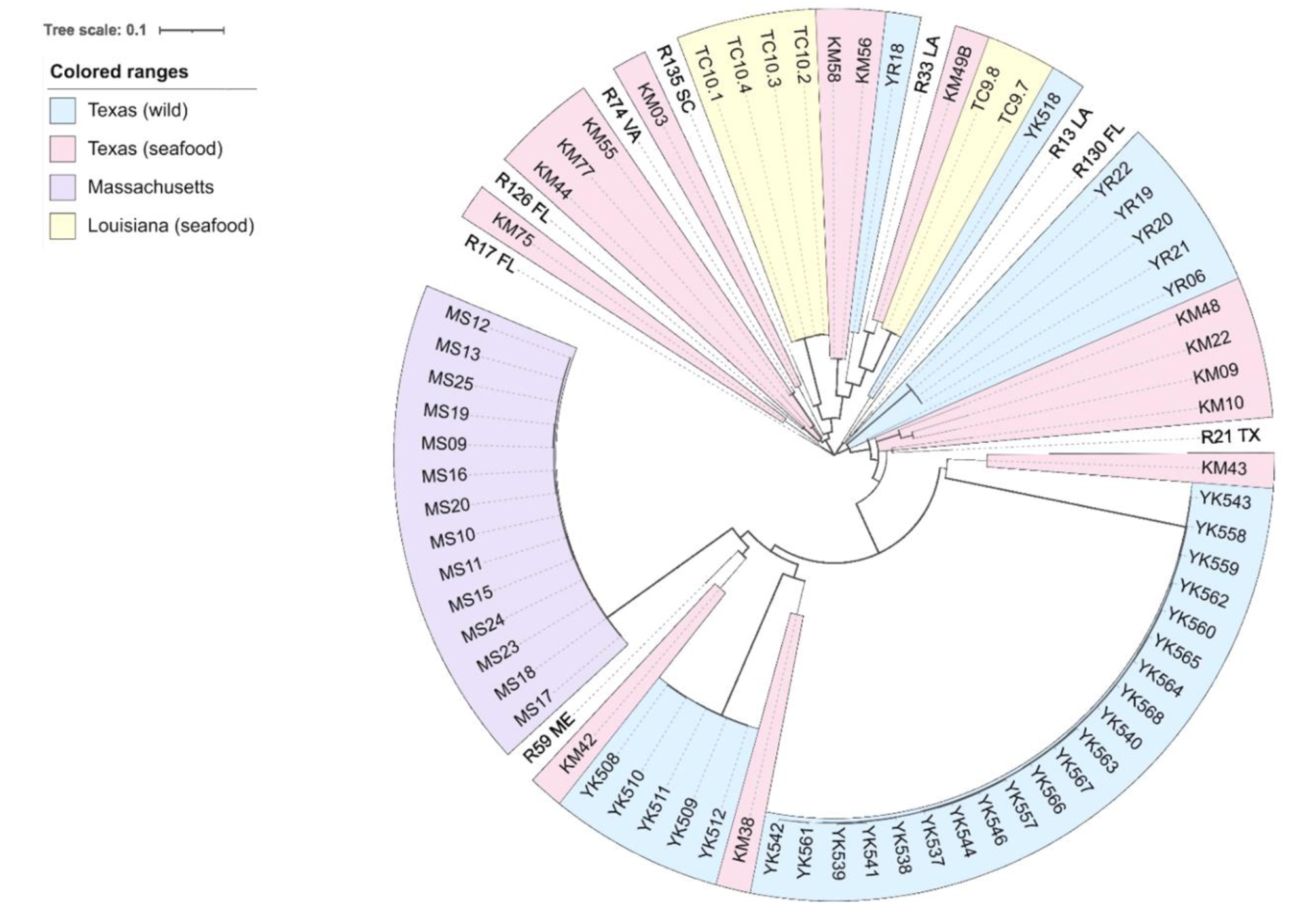
Genomic maximum likelihood tree created based on the core-genome comparison of *V. parahaemolyticus* using Roary (v3.13.0) with MAFFT (v7.490) and IQ-Tree (v2.4.0). Additional genomes of isolates collected from oysters in Florida (R17FL, R126FL, R130 FL), Louisiana (R13LA, R33LA), Maine (R59ME), South Carolina (R135SC), Texas (R21TX), and Virginia (R74VA) and originally published by Miller and colleagues (Miller et al., 2021) are incorporated into the tree for increasing phylogenetic resolution. Genomes of *V. anguillarum* (CS13 and CS14) were not included in the analysis. The tree is visualized with iTOL(v7). Colored backgrounds indicate the geographical component of the bacteria collection sources. Branch width represents bootstrap support.

The topology of the dendrogram obtained from the hierarchical cluster analysis of mass spectra was coherent; isolates formed monophyletic clades (Fig. 2). *V. parahaemolyticus* collected from Massachusetts (MS group) oysters formed a moderately robust cluster with an approximate unbiased p-value (AU) support of 92% and a bootstrap probability (BP) of 78%. Most strains isolated from wild oysters collected from Galveston Bay (YR and YK groups) also grouped into coherent clusters with varying AU and BP support values. However, isolates YK540, YK542, YK544, and YK561 clustered together with strains isolated from oysters purchased in Louisianna (TC group), rather than with other strains isolated from oysters harvested from Galveston Bay. Isolates collected from oysters purchased at the Texas seafood market (KM group), with the one exception (KM22), formed a moderately supported cluster with an AU value of 95% and a bootstrap support (BP) of 32%. However, this cluster did not constitute a well-defined clade in the phylogenetic tree. *V. anguillarum* isolates (CS13 and CS14) formed a highly robust cluster, supported by the maximum AU and BP values. The low BP value (3%) for the placement of this cluster with *V. parahaemolyticus* isolates (YK, KM) within the overall dendrogram reflects instability in global topology, a behavior commonly observed in hierarchical clustering (Shimodaira 2002).

**Figure 2.**
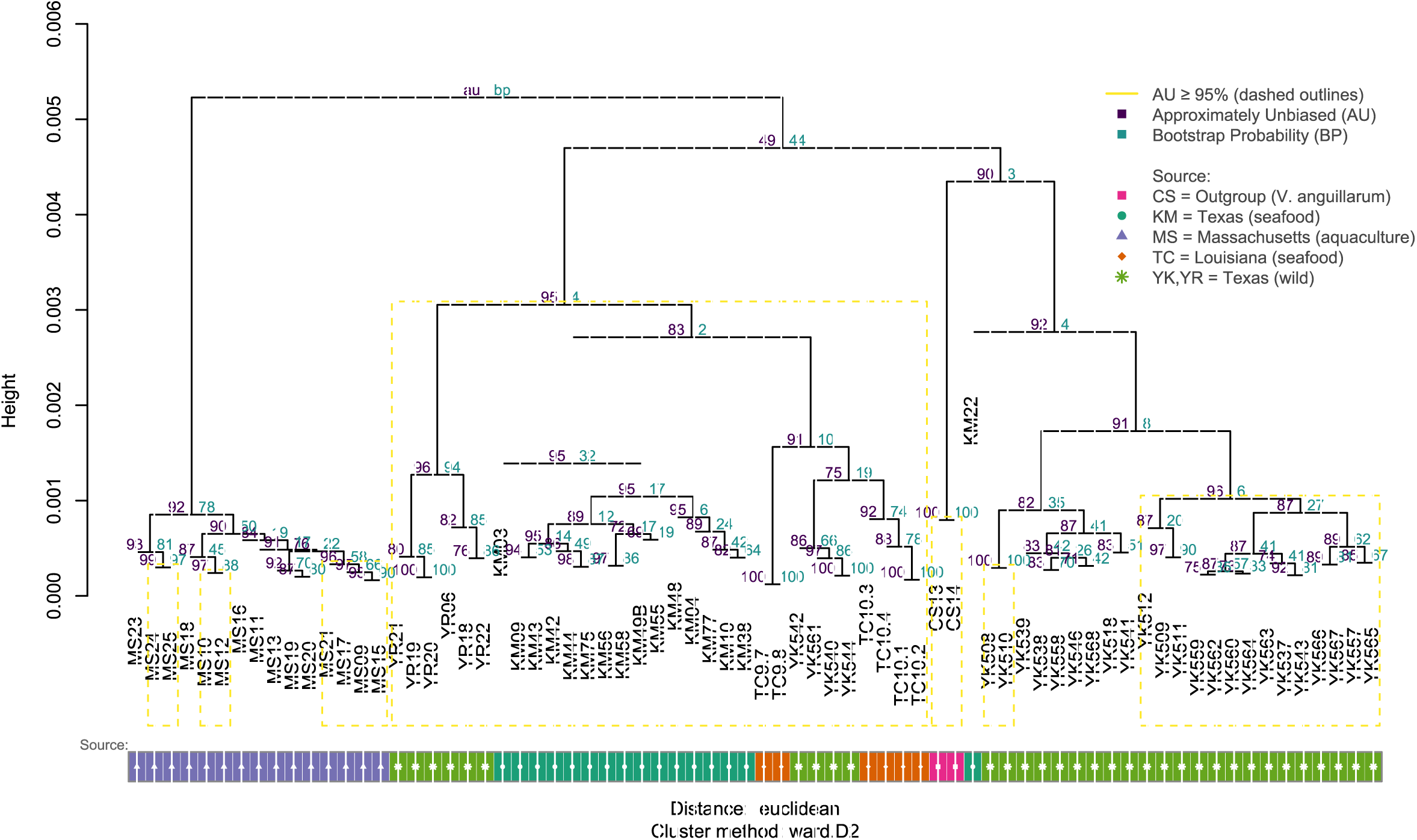
The dendrogram generated from mass spectra hierarchical cluster analysis with a custom R script illustrates protein pattern similarities among *V. parahaemolyticus* and *V. anguillarum* isolates. The approximately unbiased (AU) p-value and the bootstrap probability (BP) provide statistical support for each cluster. Clusters with AU values ≥ 95% are highlighted by PVclust (v2.2.0) with yellow edges and are considered highly reliable. The vertical height at which branches join indicates the degree of protein pattern similarity between isolates.

Pairwise comparison of isolates revealed a wider range in cosine similarity values, for comparisons of mass spectra, than the range of ANI values calculated by WGS (Fig. 3). For inter-species comparisons (*V. anguillarum* versus *V. parahaemolyticus*), cosine similarities ranged from 0.43 to 0.59 and ANI values equaled 79%. For within-species comparisons (between strains of *V. parahaemolyticus*), cosine similarities ranged from 0.61 to 0.91 and ANI values ranged from 98 to 100%.

**Figure 3.**
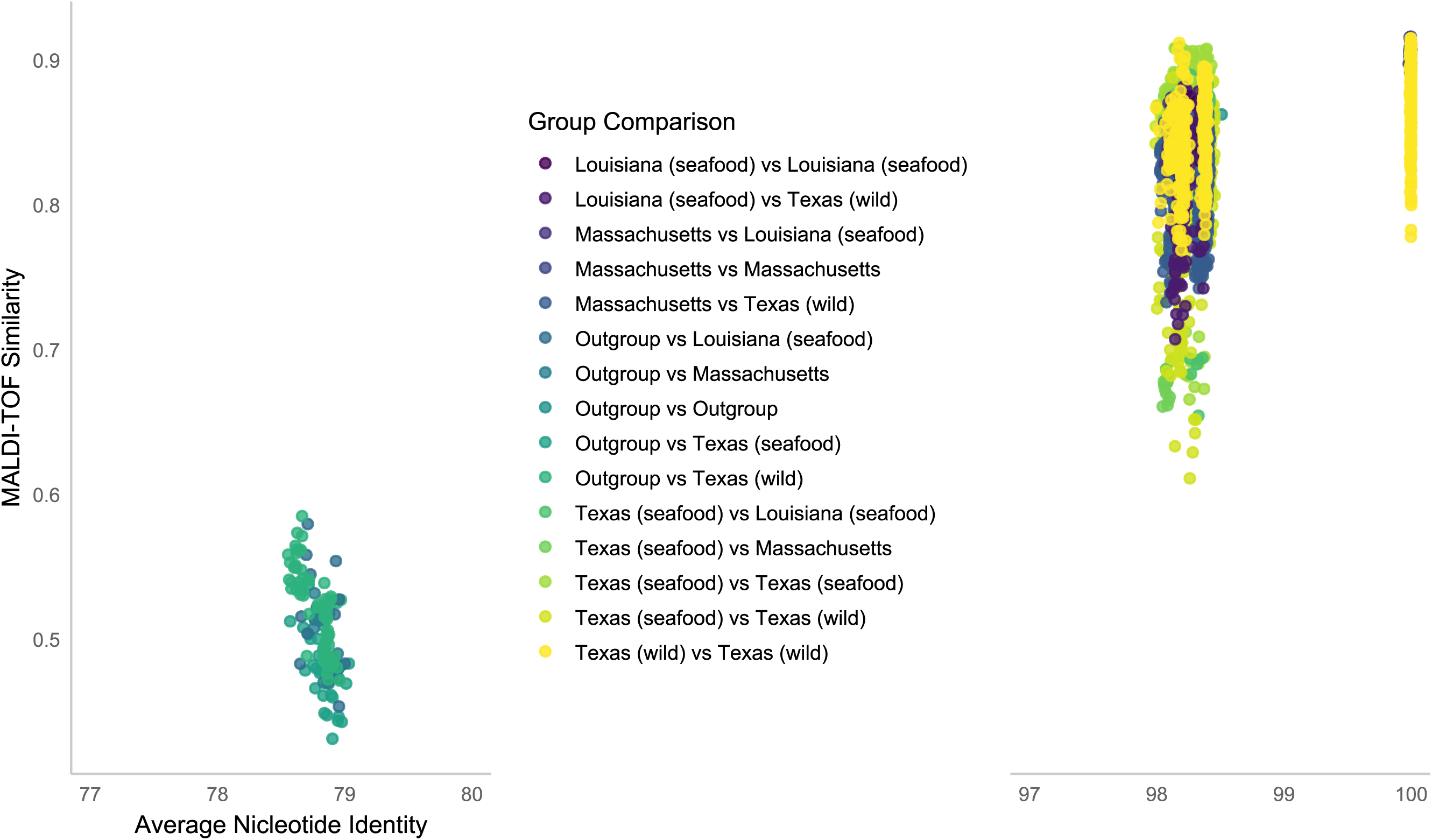
Relationship between average nucleotide identity (ANI) and MALDI-TOF similarity in pairwise comparisons of isolates. The x-axis shows ANI, and the y-axis represents the Cosine similarity coefficient derived from mass spectra pairwise comparisons. Colored dots indicate isolate pairings according to their group classifications. The left cluster, with ANI values around 79%, corresponds to comparisons between *V. parahaemolyticus* and *V. anguillarum* (Outgroup) genomes, indicating low genomic and proteomic similarity. The right cluster, with ANI values near 98%, represents comparisons among *V. parahaemolyticus* isolates, showing high genomic similarity but varying levels of MALDI-TOF spectral similarity.

Rarefaction analysis of the mass spectra showed that high spectral saturation (maximum number of distinguished proteomic features) was achieved with fewer than 50 *V. parahaemolyticus* isolates (Fig. 4). However, rarefaction analysis of COG indicated that 70 isolates were insufficient for full genomic-scale saturation (Fig. 5).

**Figure 4.**
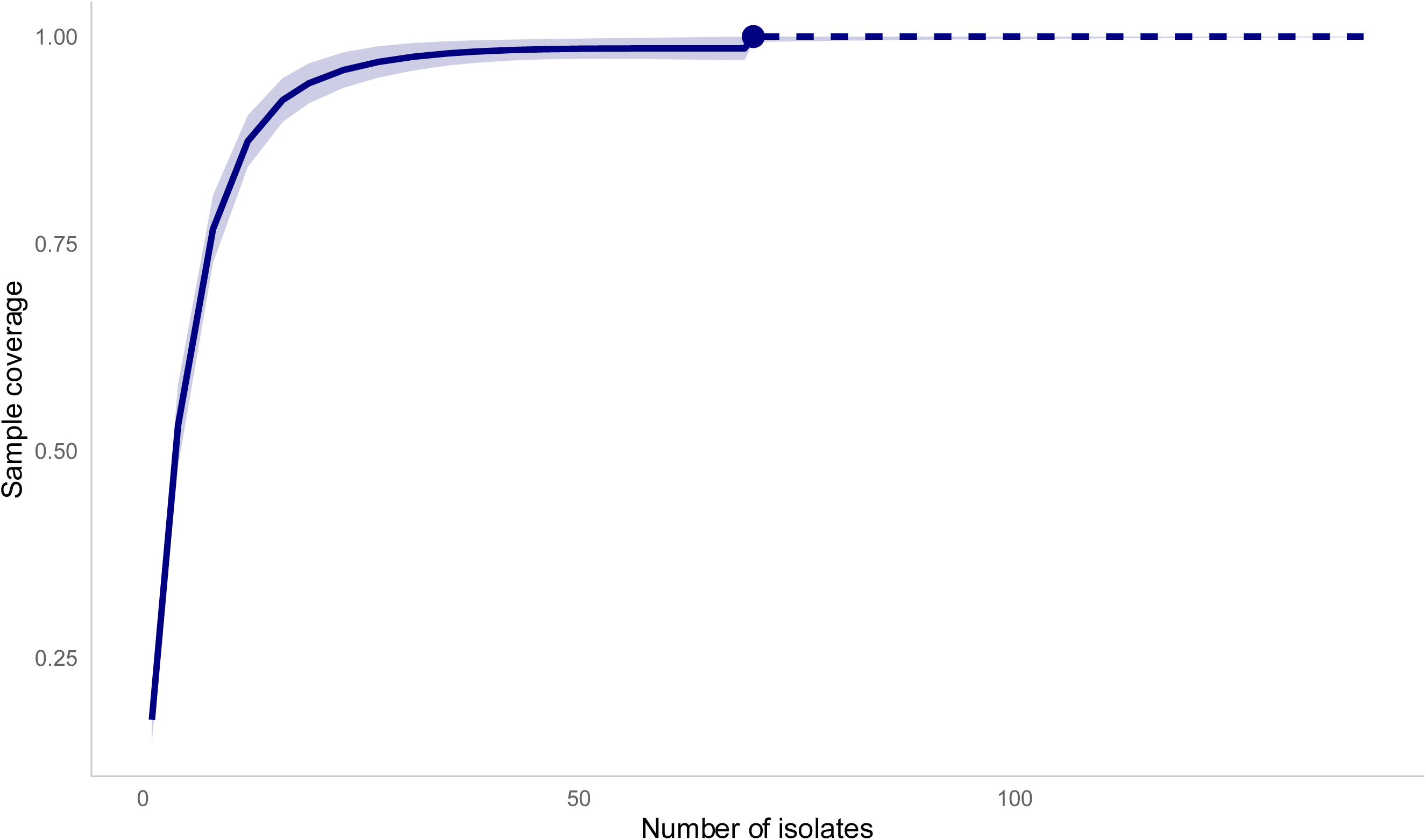
Rarefaction analysis of the MALDI-TOF spectra. The y-axis shows the estimated proportion of total proteomic diversity obtained from the analyzed spectra. The x-axis indicates the number of isolates. The solid trendline shows the observed rarefaction, and the dashed line shows extrapolated predictions with increasing numbers of isolates.

**Figure 5:**
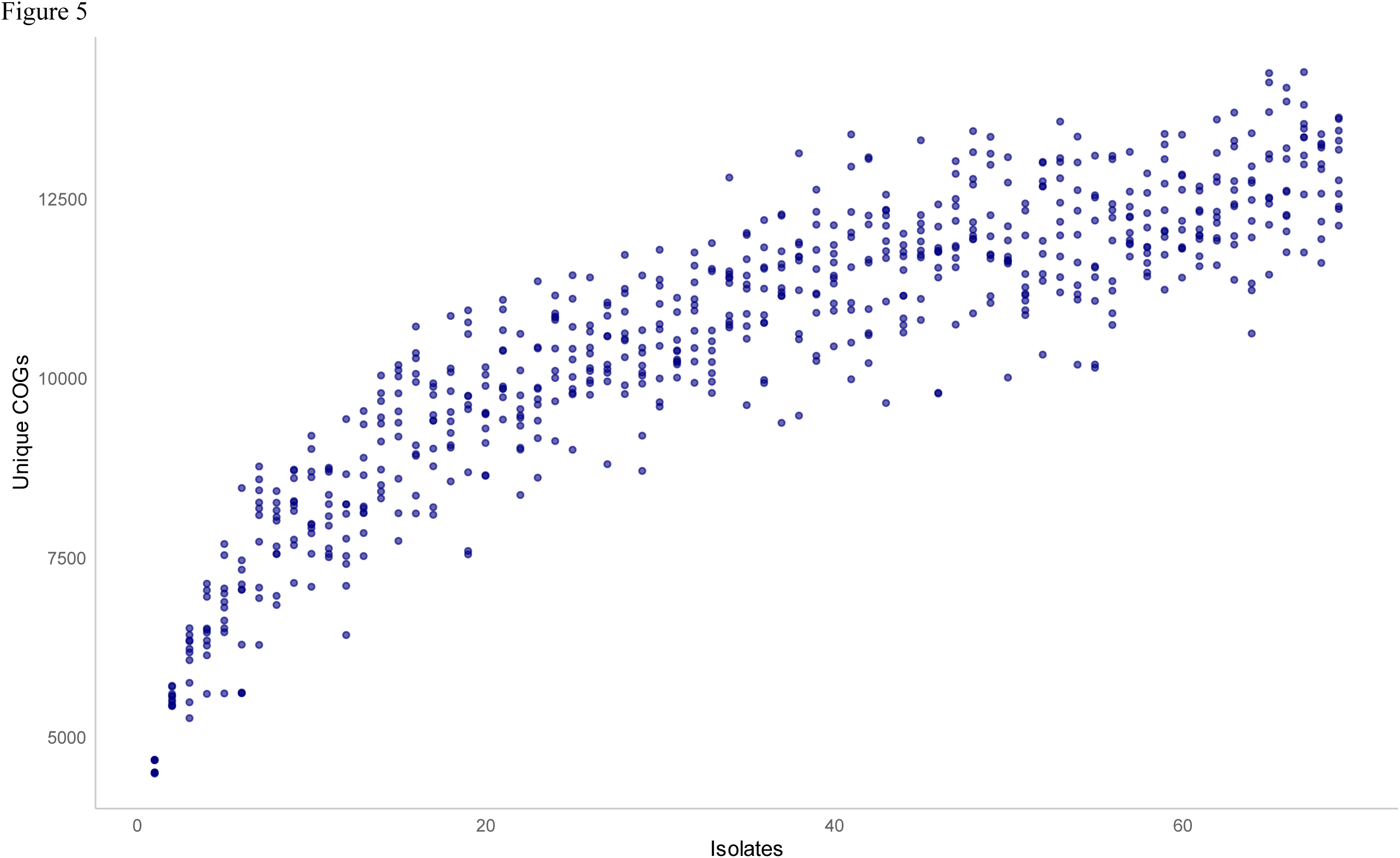
Rarefaction analysis of clusters of orthologous groups (COG). The Y-axis shows the number of COGs obtained from the gene presence/absence matrix created with Roary. Random sampling was performed 10 times for each number of isolates, and unique COGs were calculated and visualized as blue dots. The X-axis showed the number of isolates sampled.

## 4. Discussion

MALDI-TOF MS allows rapid and cost-effective identification of *V. parahaemolyticus* at the species level and provides moderate resolution for distinguishing strains. Consistent with previous studies (Hazen et al. 2009; Sulaiman et al. 2023; Vidal et al. 2020), MALDI-TOF MS effectively discriminated *V. anguillarum* from *V. parahaemolyticus*, which agreed with genomic pairwise comparisons (Fig. 3). MALDI-TOF MS showed high sensitivity (70/70) isolates for identification of *V. parahaemolyticus*, which aligns with previous reports for this species (Banerjee et al. 2025; Fasulkova et al. 2023), and supports the suitability of MALDI-TOF for species-level discrimination (Erler et al. 2015; Mougin et al. 2020). Here we report MALDI-TOF MS identified clusters of *V. parahaemolyticus* associated with particular sources, which suggests strain-level resolution. These clusters corresponded to clusters defined by WGS.

Currently, the genus *Vibrio* includes more than 150 described species (Canellas et al. 2024; Sawabe et al. 2013). Commercially available mass spectra databases, for MALDI-TOF MS systems, do not reflect the full taxonomic and genomic diversity of this group (Mougin et al. 2020). Publicly available WGS repositories contain substantially more *Vibrio* spp. genomes than are represented by MSPs included in the Bruker system. In 2015, the BioTyper database (v3.3.1.0) contained 94 MSPs for the family *Vibrionaceae*, with *V. parahaemolyticus* represented by seven MSPs (Erler et al. 2015). In the more recent version used in this study (v11.0.0.0), the database includes MSPs for 54 *Vibrio* spp, including nine MSPs of *V. parahaemolyticus* (Liu et al. 2022).

Because mass spectra-based identification primarily relies on conserved ribosomal and other highly abundant proteins (Sekiguchi et al. 2023), reliable genus- and species-level identification can be achieved despite incomplete database coverage. In contrast, strain-level resolution is more strongly influenced by database composition. In addition to the commercial database, custom MSP libraries have been developed that can improve strain-level resolution and enhance the reliability of results obtained using the Bruker MALDI Biotyper (Erler et al. 2015; Liu et al. 2022; Moussa Pouly et al.). In the present study, *V. parahaemolyticus* isolates were identified exclusively using the standard commercial MALDI Biotyper database to avoid compatibility issues associated with the use of multiple databases and to ensure methodological consistency and standardization. Even with this limited database, 70 of 70 of the isolates were identified, with high confidence (scores 2.3 – 2.6, File S1) and complete agreement between species identification between MALDI-TOF MS and WGS. Isolates that were not identified as *V. parahaemolyticus,* with the Bruker MALDI Biotyper system, were not further analyzed in this study.

Despite overlapping cosine similarity values within species, MALDI-TOF MS captured subtle proteomic differences between isolates, suggesting source-dependent variation. Hierarchical clustering of spectra revealed consistent grouping of isolates depending upon the source; however, clusters consisting of bacteria isolated from oysters purchased at seafood markets had low bootstrap support. Indeed, genetically indistinguishable isolates, KM03 and KM04, showed clearly different mass-spectra profiles. This may reflect a higher degree of phenotypic plasticity due to differences in storage conditions and environmental stress factors in seafood markets (Audemard et al. 2022). Strains isolated from oysters collected from an aquaculture facility in Massachusetts formed a coherent clade that corresponded to clusters defined by WGS. Aquaculture conditions may select for specific bacterial genotypes and, as a result, this clade may reflect local adaptation (Bosi et al. 2024; Kopprio et al. 2017); however, only one set of oysters was collected from Massachusetts in this study. This limits broader interpretation of the data.

Mass spectra reported herein were all generated with one MALDI-TOF MS system by one group. However, in terms of bacterial identification, MALDI-TOF MS systems from different manufacturers give comparable results (Cherkaoui et al. 2010; Dichtl et al. 2023; Sibińska et al. 2024) and MALDI-TOF MS results are generally reproducible between laboratories (Mellmann et al. 2009). Mass spectra generated from bacteria by MALDI-TOF MS may be sensitive to media formulation and culture conditions (Balážová et al. 2014; Cuénod et al. 2021; Haider et al. 2023). These factors, and sample extraction protocol (Goldstein et al. 2013), can influence the quality of mass spectra generated and the confidence of bacterial identification by MALDI-TOF MS systems (Topić Popović et al. 2023). For example, sporulation by Bacillus species, which increases rapidly as colonies age, can interfere with identification by MALDI-TOF MS (Baek et al. 2025).

*Vibrio* species, because of their fast growth rate, appear particularly sensitive to colony age, when mass spectra are generated by “on target” extraction (Kazazić et al. 2019). All isolates for this research were cultured under identical conditions, including the same media, incubation time, and temperature, and only the standard full extraction protocol, recommended by Bruker, was used in this study. This “in-tube” protocol improves identification of bacteria (Alatoom et al. 2011) and biosafety (Rudrik et al. 2017) relative to on target protocols and provides near-perfect specificity and sensitivity for identification of *V. parahaemolyticus* (Banerjee et al. 2025). Preparation of *Vibrio* species by extraction with trifluoroacetic acid from liquid cultures for MALDI-TOF MS shows promise (Cho et al. 2017); however, the quality of mass spectra generated by MALDI-TOF MS from liquid cultures is more sensitive, than colonies on agar plates, to the growth stage of the cells (Vargha et al. 2006).

Pan-genome analysis of 70 *V. parahaemolyticus* isolates identified 14,867 genes, of which 2,686 (approximately 18%) were classified as essential genes because they were present in ≥ 99% of isolates. This degree of genetic conservation is consistent with the fundamental principles of microbial identification by MALDI-TOF MS, which is predominantly based on mass spectra patterns consisting of distinct peaks of ionized conserved proteins in the 2- to 15-kDa range (Sekiguchi et al. 2023). Due to these conserved biological markers, MALDI-TOF MS is an effective tool for *V. parahaemolyticus* identification; however, it is inferior to genetic methods in differentiating bacteria at the strain level. This limitation should be considered when working with epidemiologically significant isolates (Li et al. 2018; Rahmani et al. 2021).

AMR profiles of the *V. parahaemolyticus* analyzed herein did not correspond to clusters defined by mass spectra generated by MALDI-TOF MS. All isolates of this species contained tetracycline resistance genes and were heterogeneous in the distribution of β-lactam resistance genes. These differences were not reflected in the cluster dendrogram, indicating that MALDI-TOF MS could not differentiate antibiotic-resistant and -sensitive strains of *V. parahaemolyticus.* This is consistent with previous reports (Hazen et al. 2009) and suggests MALDI-TOF MS should be combined with WGS or other methods for AMR profiling (Li et al. 2018).

Rarefaction analysis of the mass spectra demonstrated high spectral saturation (the maximum number of distinguishable proteomic features) was reached with fewer than 50 *V. parahaemolyticus* isolates (Fig. 4). In contrast, rarefaction analysis of COGs indicated that even 70 isolates did not reach saturation at the genomic level (Fig. 5). This suggests that the WGS has a higher resolution for typing strains of *V. parahaemolyticus* than MALDI-TOF MS. MALDI-TOF MS primarily targets ribosomal proteins, whose composition varies among bacterial species (Cheng et al. 2018; Clark et al. 2013; Suarez et al. 2013). WGS, by definition, describes the entire genome and remains the preferred approach for high-precision strain differentiation (Banerjee et al. 2025; Mazhari et al. 2025). A hybrid strategy, using MALDI-TOF MS for high-throughput screening followed by WGS for representative isolates (Rahmani et al. 2021), may provide an optimal balance between resolution, speed and cost.

## Conclusions

MALDI-TOF MS can be used to identify strains of *V. parahaemolyticus* isolated from oysters. This proteomic method effectively distinguished *V. parahaemolyticus* strains isolated from oysters harvested from different locations and allowed attribution of the source of the isolates with a resolution comparable to, but not as high as, WGS.

## Acknowledgements

Authors thank Kimberly Jeffrey and the team of the Texas Parks and Wildlife Department Marine Resources Laboratory (Dickinson, TX) for providing oysters and Taylor Baccus, Kirsten Lenz, Emily Pechota, Maria Borunda, Mina Chhikara, Mario Munoz, Meghana Budankayala, and Aspen Earle for preliminary results. This research was supported by the National Aeronautics and Space Administration (NASA) grant (23-BPSF23-0008), Environmental Protection Agency (EPA) grant (02D18322) and National Science Foundation (NSF) grant (2320765) to MGL. Support was also received from the Food and Drug Administration (FDA) Laboratory Flexible Funding Model (LFFM) cooperative agreement for whole genome sequencing at the Texas Public Health Laboratory. The contents are those of the author(s) and do not necessarily represent the official views of, nor an endorsement, by FDA/HHS, or the U.S. Government.

## Supplementary Material and Data Availability

A custom MSP library (.btmsp file) of *V. parahaemolyticus* and the raw MALDI-TOF MS spectra used for bacterial identification are available on Zenodo (https://zenodo.org/records/17188941). Genomic sequences and assemblies are available in Bioprojects PRJNA266293 and PRJNA1313351.

## Supplemental Figure Legends

**File S1.** Metadata and basic statistics for *V. parahaemolyticus* isolates

**File S2.** Presence/absence heatmap of AMR and virulence-associated genes across *V. parahaemolyticus* isolates. Each row represents an isolate, and each column corresponds to an individual gene. Orange boxes indicate the presence of AMR genes, blue boxes indicate the presence of virulence-associated genes, and white boxes indicate the absence of the gene. AMR genes include β-lactam resistance (ampC, blaCARB variants), trimethoprim resistance (dfrA), and tetracycline resistance genes (tet(34) and tet(35)). Virulence genes include regulators and accessory factors (exsA, exsD, sycN, tyeA, tlh, vat, virG, vecA, vxsC) as well as components of the Type III secretion system (T3SS), including regulatory genes (vcr family), effector proteins (vop family), and structural genes (vsc family).

**File S3.** Basic statistics for *V. anguillarum* hybrid genome assembly and annotation.

**File S4.** Presence/absence heatmap of AMR and virulence-associated genes across *V. anguillarum* isolates CS13 and CS14. Rows represent isolates, and columns represent individual genes. Orange boxes correspond to AMR genes, blue boxes correspond to virulence-associated genes, white boxes indicate the absence of the gene. AMR profile includes vancomycin resistance (vanT). Virulence genes are organized into functional groups, including regulators (CRP, rsmA), motility and chemotaxis genes (cheW, cheY, flaA, flg and fli families), toxin and virulence-associated systems (vab and vop families), iron acquisition systems (hmu family), and other virulence-associated factors (RbmA, RbmC, cpsF, dthP, epsG, hcp-2, hthH, vah1, vibriolysin).

**File S5.** Circular representation of genetic content distribution on chromosome 1 of *V. anguillarum* isolate CS13. The size of the chromosome is 2.6 Mbp. Rings represent CDS on the forward and reverse strands (light grey bars) and annotated RNA genes: transfer RNA (green bars), ribosomal RNA (red bars), non-coding RNAs (orange bars), and regulatory non-coding RNA (light blue bars). CRISPR arrays are represented by lilac bars in the legend; however, no CRISPR elements were annotated in the chromosome (middle ring). GC content variation is shown as a separate blue-and-orange circle that includes the position of the origin of replication (oriC). GC skew is displayed as a green-and-purple circle. The outer two rings indicate functional annotation of protein-coding genes according to COG categories, represented by different colors: RNA processing and modification (A, dark red), chromatin structure and dynamics (B, purple), energy production and conversion (C, light violet), cell cycle control and chromosome partitioning (D, orange), amino acid transport and metabolism (E, pale yellow), nucleotide transport and metabolism (F, red), carbohydrate transport and metabolism (G, salmon), coenzyme transport and metabolism (H, light green), lipid transport and metabolism (I, very light green), translation and ribosomal biogenesis (J, light blue), transcription (K, yellow-green), replication, recombination and repair (L, blue), cell wall/membrane/envelope biogenesis (M, lavender), cell motility (N, pale purple), post-translational modification and chaperones (O, green), inorganic ion transport and metabolism (P, pink), secondary metabolite biosynthesis and transport (Q, dark blue), general function prediction only (R, turquoise), function unknown (S, black), signal transduction mechanisms (T, light orange), intracellular trafficking and secretion (U, peach), defense mechanisms (V, light red), mobilome elements including prophages and transposons (X, grey).

**File S6.** Circular representation of genetic content distribution on chromosome I of *V. anguillarum* isolate CS13. The size of the chromosome is 1.2 Mbp. Rings represent CDS on the forward and reverse strands (light grey bars) and annotated RNA genes: transfer RNA (green bars), ribosomal RNA (red bars), non-coding RNAs (orange bars), and regulatory non-coding RNA (light blue bars). CRISPR arrays are represented by lilac bars in the legend; however, no CRISPR elements were annotated in the chromosome (middle ring). GC content variation is shown as a separate blue-and-orange circle that includes the position of the origin of replication (oriC). GC skew is displayed as a green-and-purple circle. The outer two rings indicate functional annotation of protein-coding genes according to COG categories, represented by different colors: RNA processing and modification (A, dark red), chromatin structure and dynamics (B, purple), energy production and conversion (C, light violet), cell cycle control and chromosome partitioning (D, orange), amino acid transport and metabolism (E, pale yellow), nucleotide transport and metabolism (F, red), carbohydrate transport and metabolism (G, salmon), coenzyme transport and metabolism (H, light green), lipid transport and metabolism (I, very light green), translation and ribosomal biogenesis (J, light blue), transcription (K, yellow-green), replication, recombination and repair (L, blue), cell wall/membrane/envelope biogenesis (M, lavender), cell motility (N, pale purple), post-translational modification and chaperones (O, green), inorganic ion transport and metabolism (P, pink), secondary metabolite biosynthesis and transport (Q, dark blue), general function prediction only (R, turquoise), function unknown (S, black), signal transduction mechanisms (T, light orange), intracellular trafficking and secretion (U, peach), defense mechanisms (V, light red), mobilome elements including prophages and transposons (X, grey).

**File S7.** Circular representation of genetic content distribution on chromosome 1 of *V. anguillarum* isolate CS14. The size of the chromosome is 2.8 Mbp. Rings represent CDS on the forward and reverse strands (light grey bars) and annotated RNA genes: transfer RNA (green bars), ribosomal RNA (red bars), non-coding RNAs (orange bars), and regulatory non-coding RNA (light blue bars). CRISPR arrays are represented by lilac bars in the legend; however, no CRISPR elements were annotated in the chromosome (middle ring). GC content variation is shown as a separate blue-and-orange circle that includes the position of the origin of replication (oriC). GC skew is displayed as a green-and-purple circle. The outer two rings indicate functional annotation of protein-coding genes according to COG categories, represented by different colors: RNA processing and modification (A, dark red), chromatin structure and dynamics (B, purple), energy production and conversion (C, light violet), cell cycle control and chromosome partitioning (D, orange), amino acid transport and metabolism (E, pale yellow), nucleotide transport and metabolism (F, red), carbohydrate transport and metabolism (G, salmon), coenzyme transport and metabolism (H, light green), lipid transport and metabolism (I, very light green), translation and ribosomal biogenesis (J, light blue), transcription (K, yellow-green), replication, recombination and repair (L, blue), cell wall/membrane/envelope biogenesis (M, lavender), cell motility (N, pale purple), post-translational modification and chaperones (O, green), inorganic ion transport and metabolism (P, pink), secondary metabolite biosynthesis and transport (Q, dark blue), general function prediction only (R, turquoise), function unknown (S, black), signal transduction mechanisms (T, light orange), intracellular trafficking and secretion (U, peach), defense mechanisms (V, light red), mobilome elements including prophages and transposons (X, grey).

**File S8.** Circular representation of genetic content distribution on chromosome I of *V. anguillarum* isolate CS14. The size of the chromosome is 1.1 Mbp. Rings represent CDS on the forward and reverse strands (light grey bars) and annotated RNA genes: transfer RNA (green bars), ribosomal RNA (red bars), non-coding RNAs (orange bars), and regulatory non-coding RNA (light blue bars). CRISPR arrays are represented by lilac bars in the legend; however, no CRISPR elements were annotated in the chromosome (middle ring). GC content variation is shown as a separate blue-and-orange circle that includes the position of the origin of replication (oriC). GC skew is displayed as a green-and-purple circle. The outer two rings indicate functional annotation of protein-coding genes according to COG categories, represented by different colors: RNA processing and modification (A, dark red), chromatin structure and dynamics (B, purple), energy production and conversion (C, light violet), cell cycle control and chromosome partitioning (D, orange), amino acid transport and metabolism (E, pale yellow), nucleotide transport and metabolism (F, red), carbohydrate transport and metabolism (G, salmon), coenzyme transport and metabolism (H, light green), lipid transport and metabolism (I, very light green), translation and ribosomal biogenesis (J, light blue), transcription (K, yellow-green), replication, recombination and repair (L, blue), cell wall/membrane/envelope biogenesis (M, lavender), cell motility (N, pale purple), post-translational modification and chaperones (O, green), inorganic ion transport and metabolism (P, pink), secondary metabolite biosynthesis and transport (Q, dark blue), general function prediction only (R, turquoise), function unknown (S, black), signal transduction mechanisms (T, light orange), intracellular trafficking and secretion (U, peach), defense mechanisms (V, light red), mobilome elements including prophages and transposons (X, grey).

## Abbreviations

ANI: average nucleotide identity
AMR: antimicrobial resistance
AU: approximate unbiased
BP: bootstrap probability
CDS: protein-coding sequences
COG: cluster of orthologous groups
MA: marine agar
MALDI-TOF MS: matrix-assisted laser desorption/ionization time-of-flight mass spectrometry
MSP: main spectrum profile
MTUs: MALDI-TOF taxonomic units
RGI: resistance gene identifier
SNR: signal-to-noise ratio
VF: virulence factor
VFDB: virulence factor database
WGS: whole genome sequencing

## References

1. Alatoom AA, Cunningham SA, Ihde SM, Mandrekar J, Patel R (2011) Comparison of direct colony method versus extraction method for identification of Gram-positive cocci by use of Bruker Biotyper matrix-assisted laser desorption ionization–time of flight mass spectrometry. J Clin Microbiol 49:2868–2873. doi:10.1128/jcm.00506-11.

2. Alcock BP, Huynh W, Chalil R, Smith KW, Raphenya Amogelang R, Wlodarski MA, Edalatmand A, Petkau A, Syed SA, Tsang KK, Baker SJC, Dave M, McCarthy Madeline C, Mukiri KM, Nasir JA, Golbon B, Imtiaz H, Jiang X, Kaur K, Kwong M, Liang ZC, Niu KC, Shan P, Yang JYJ, Gray Kristen L, Hoad GR, Jia B, Bhando T, Carfrae Lindsey A, Farha Maya A, French S, Gordzevich R, Rachwalski K, Tu Megan M, Bordeleau E, Dooley D, Griffiths E, Zubyk HL, Brown ED, Maguire F, Beiko Robert G, Hsiao WWL, Brinkman FSL, Van Domselaar G, McArthur AG (2023) CARD 2023: expanded curation, support for machine learning, and resistome prediction at the Comprehensive Antibiotic Resistance Database. Nucleic Acids Res 51:D690–D699. 10.1093/nar/gkac920.

3. Alipour M, Issazadeh K, Soleimani J (2014) Isolation and identification of *Vibrio parahaemolyticus* from seawater and sediment samples in the southern coast of the Caspian Sea. Comp Clin Path 23:129–133. 10.1007/s00580-012-1583-6.

4. Audemard C, Ben-Horin T, Kator HI, Reece KS (2022) *Vibrio vulnificus* and *Vibrio parahaemolyticus* in oysters under low tidal range conditions: is seawater analysis useful for risk assessment? Foods 11:4065. https://www.mdpi.com/2304-8158/11/24/4065

5. Baek B, Park YH, Jeon J-M, Shim H-Y, Lee E-K, Hong M-J, Bae Y-W, An J-H, Shin I-C, Jung HS (2025) Optimizing MALDI-TOF mass spectrometry for the identification of *Bacillus cereus*: the impact of sporulation and cultivation time. Int J Molec Sci 26:4355. https://www.mdpi.com/1422-0067/26/9/4355

6. Baker-Austin C, Trinanes JA, Taylor NGH, Hartnell R, Siitonen A, Martinez-Urtaza J (2013) Emerging *Vibrio* risk at high latitudes in response to ocean warming. Nature Clim Change 3:73–77. 10.1038/nclimate1628.

7. Balážová T, Makovcová J, Šedo O, Slaný M, Faldyna M, Zdráhal Z (2014) The influence of culture conditions on the identification of *Mycobacterium* species by MALDI-TOF MS profiling. FEMS Microbiol Lett 353:77–84. 10.1111/1574-6968.12408.

8. Banerjee S, Flint A, Brosseau MB, Weedmark K, Shutinoski B (2025) Evaluation of MALDI-TOF for identification of *Vibrio cholerae* and *Vibrio parahaemolyticus* from growth on agar media. Appl Microbiol Biotechnol 109:5. 10.1007/s00253-024-13385-y.

9. Benson G (1999) Tandem repeats finder: a program to analyze DNA sequences. Nucleic Acids Res 27:573–580. 10.1093/nar/27.2.573.

10. Besemer J, Lomsadze A, Borodovsky M (2001) GeneMarkS: a self-training method for prediction of gene starts in microbial genomes. Implications for finding sequence motifs in regulatory regions. Nucleic Acids Res 29:2607–2618. 10.1093/nar/29.12.2607.

11. Boonstra M, Fouz B, van Gelderen B, Dalsgaard I, Madsen L, Jansson E, Amaro C, Haenen O (2023) Fast and accurate identification by MALDI-TOF of the zoonotic serovar E of *Vibrio vulnificus* linked to eel culture. J Fish Dis 46:445–452. 10.1111/jfd.13756.

12. Bosi E, Taviani E, Avesani A, Doni L, Auguste M, Oliveri C, Leonessi M, Martinez-Urtaza J, Vetriani C, Vezzulli L (2024) Pan-genome provides insights into *Vibrio* evolution and adaptation to deep-sea hydrothermal vents. Genome Biol Evol 1610.1093/gbe/evae131.

13. Brown B, Allard M, Bazaco MC, Blankenship J, Minor T (2021) An economic evaluation of the whole genome sequencing source tracking program in the U.S. PLoS One 16:e0258262. doi: 10.1371/journal.pone.0258262.

14. Canellas ALB, Faria AR, Dias GR, Teixeira LM, Laport MS (2024) Polyphasic identification of *Vibrio* species from aquatic sources using mass spectrometry, housekeeping gene sequencing and whole genome analysis. Sci Rep 14:26250. 10.1038/s41598-024-77919-0.

15. Chan PP, Lowe TM, tRNAscan-SE: Searching for tRNA Genes in Genomic Sequences. in: M. Kollmar, (Ed.), Gene Prediction: Methods and Protocols, Springer New York, New York, NY, 2019, pp. 1–14.

16. Chen S (2023) Ultrafast one-pass FASTQ data preprocessing, quality control, and deduplication using fastp. iMeta 2:e107. 10.1002/imt2.107.

17. Cheng D, Qiao L, Horvatovich P (2018) Toward spectral library-free matrix-assisted laser desorption/ionization time-of-flight mass spectrometry bacterial identification. J Proteome Res 17:2124–2130. 10.1021/acs.jproteome.8b00065.

18. Cherkaoui A, Hibbs J, Emonet S, Tangomo M, Girard M, Francois P, Schrenzel J (2010) Comparison of two matrix-assisted laser desorption ionization-time of flight mass spectrometry methods with conventional phenotypic identification for routine identification of bacteria to the species level. J Clin Microbiol 48:1169–1175. doi:10.1128/jcm.01881-09.

19. Cho Y, Kim E, Han SK, Yang SM, Kim MJ, Kim HJ, Kim CG, Choo DW, Kim YR, Kim HY (2017) Rapid identification of *Vibrio* species isolated from the southern coastal regions of Korea by MALDI-TOF Mass Spectrometry and comparison of MALDI sample preparation methods. J Microbiol Biotechnol 27:1593–1601. 10.4014/jmb.1704.04056.

20. Clark AE, Kaleta EJ, Arora A, Wolk DM (2013) Matrix-assisted laser desorption ionization–time of flight mass spectrometry: a fundamental shift in the routine practice of clinical microbiology. Clin Microbiol Rev 26:547–603. doi:10.1128/cmr.00072-12.

21. Couderc C, Nappez C, Drancourt M (2012) Comparing inactivation protocols of *Yersinia* organisms for identification with matrix-assisted laser desorption/ionization time-of-flight mass spectrometry. Rapid Commun Mass Spectrom 26:710–714. 10.1002/rcm.6152.

22. Cuénod A, Foucault F, Pflüger V, Egli A (2021) Factors associated with MALDI-TOF mass spectral quality of species identification in clinical routine diagnostics. Front Cell Infect Microbiol 11:646648. 10.3389/fcimb.2021.646648.

23. Daniels NA, MacKinnon L, Bishop R, Altekruse S, Ray B, Hammond RM, Thompson S, Wilson S, Bean NH, Griffin PM, Slutsker L (2000) *Vibrio parahaemolyticus* infections in the United States, 1973–1998. J Infect Dis 181:1661–1666. 10.1086/315459.

24. Davis JW, Sizemore RK (1982) Incidence of *Vibrio* species associated with blue crabs (*Callinectes sapidus*) collected from Galveston Bay, Texas. Appl Envir Microbiol 43:1092–7. 10.1128/aem.43.5.1092-1097.1982.

25. de Jong E, de Jong AS, Smidts-van den Berg N, Rentenaar RJ (2013) Differentiation of *Raoultella ornithinolytica/planticola* and *Klebsiella oxytoca* clinical isolates by matrix-assisted laser desorption/ionization-time of flight mass spectrometry. Diagn Microbiol Infect Dis 75:431–433. 10.1016/j.diagmicrobio.2012.12.009.

26. Dichtl K, Klugherz I, Greimel H, Luxner J, Köberl J, Friedl S, Steinmetz I, Leitner E (2023) A head-to-head comparison of three MALDI-TOF mass spectrometry systems with 16S rRNA gene sequencing. J Clin Microbiol 61:e0191322. 10.1128/jcm.01913-22.

27. Donohue MJ, Smallwood AW, Pfaller S, Rodgers M, Shoemaker JA (2006) The development of a matrix-assisted laser desorption/ionization mass spectrometry-based method for the protein fingerprinting and identification of *Aeromonas* species using whole cells. J Microbiol Method 65:380–389. 10.1016/j.mimet.2005.08.005.

28. Erler R, Wichels A, Heinemeyer E-A, Hauk G, Hippelein M, Reyes NT, Gerdts G (2015) VibrioBase: A MALDI-TOF MS database for fast identification of *Vibrio* spp. that are potentially pathogenic in humans. Syst Appl Microbiol 38:16–25. doi: 10.1016/j.syapm.2014.10.009.

29. Fasulkova R, Orozova P, Stratev D (2023) Identification of *Vibrio parahemolyticus* isolated from seafood via matrix-assisted laser desorption/ionization time of flight mass spectrometry. J Food Qual Haz Control 10:135–141. https://publish.kne-publishing.com/index.php/JFQHC/article/view/13644

30. Flynn JM, Hubley R, Goubert C, Rosen J, Clark AG, Feschotte C, Smit AF (2020) RepeatModeler2 for automated genomic discovery of transposable element families. PNAS 117:9451–9457. doi:10.1073/pnas.1921046117.

31. Frans I, Michiels CW, Bossier P, Willems KA, Lievens B, Rediers H (2011) *Vibrio anguillarum* as a fish pathogen: virulence factors, diagnosis and prevention. J Fish Dis 34:643–661. 10.1111/j.1365-2761.2011.01279.x.

32. Galán JE, Collmer A (1999) Type III secretion machines: bacterial devices for protein delivery into host cells. Science 284:1322–1328. doi:10.1126/science.284.5418.1322.

33. Gibb S, Strimmer K (2012) MALDIquant: a versatile R package for the analysis of mass spectrometry data. Bioinformatics 28:2270–2271. doi: 10.1093/bioinformatics/bts447.

34. Gode-Potratz CJ, McCarter LL (2011) Quorum sensing and silencing in *Vibrio parahaemolyticus*. J Bacteriol 193:4224–4237. doi:10.1128/jb.00432-11.

35. Goldstein JE, Zhang L, Borror CM, Rago JV, Sandrin TR (2013) Culture conditions and sample preparation methods affect spectrum quality and reproducibility during profiling of *Staphylococcus aureus* with matrix-assisted laser desorption/ionization time-of-flight mass spectrometry. Lett Appl Microbiol 57:144–150. 10.1111/lam.12092.

36. Gomes E, Araújo D, Nogueira T, Oliveira R, Silva S, Oliveira LVN, Azevedo NF, Almeida C, Castro J (2025) Advances in whole genome sequencing for foodborne pathogens: implications for clinical infectious disease surveillance and public health. Front Cell Infect Microbiol 15:1593219. 10.3389/fcimb.2025.1593219.

37. Haider A, Ringer M, Kotroczó Z, Mohácsi-Farkas C, Kocsis T (2023) The current level of MALDI-TOF MS applications in the detection of microorganisms: a short review of benefits and limitations. Microbiol Res 14:80–90. https://www.mdpi.com/2036-7481/14/1/8

38. Hazen TH, Martinez RJ, Chen Y, Lafon PC, Garrett NM, Parsons MB, Bopp CA, Sullards MC, Sobecky PA (2009) Rapid identification of *Vibrio parahaemolyticus* by whole-cell matrix-assisted laser desorption ionization-time of flight mass spectrometry. Appl Envir Microbiol 75:6745–6756. doi:10.1128/AEM.01171-09.

39. Hornik K, Buchta C, Zeileis A (2009) Open-source machine learning: R meets Weka. Comput Stat 24:225–232. doi: 10.1007/s00180-008-0119-7.

40. Hyatt D, Chen G-L, LoCascio PF, Land ML, Larimer FW, Hauser LJ (2010) Prodigal: prokaryotic gene recognition and translation initiation site identification. BMC Bioinform 11:119. 10.1186/1471-2105-11-119.

41. Jain C, Rodriguez-R LM, Phillippy AM, Konstantinidis KT, Aluru S (2018) High throughput ANI analysis of 90K prokaryotic genomes reveals clear species boundaries. Nat Commun 9:5114. 10.1038/s41467-018-07641-9.

42. Kaysner CA, DePaola A, Jones J, *Vibrio*, Bacteriological Analytical Manual, FDA, 2004, pp. 1–45.

43. Kazazić SP, Topić Popović N, Strunjak-Perović I, Babić S, Florio D, Fioravanti M, Bojanić K, Čož-Rakovac R (2019) Matrix-assisted laser desorption/ionization time of flight mass spectrometry identification of *Vibrio* (*Listonella*) *anguillarum* isolated from sea bass and sea bream. PLOS ONE 14:e0225343. 10.1371/journal.pone.0225343.

44. Kopprio GA, Streitenberger ME, Okuno K, Baldini M, Biancalana F, Fricke A, Martínez A, Neogi SB, Koch BP, Yamasaki S, Lara RJ (2017) Biogeochemical and hydrological drivers of the dynamics of *Vibrio* species in two Patagonian estuaries. Sci Total Environ 579:646–656. 10.1016/j.scitotenv.2016.11.045.

45. LaMontagne MG, Tran PL, Benavidez A, Morano LD (2021) Development of an inexpensive matrix-assisted laser desorption—time of flight mass spectrometry method for the identification of endophytes and rhizobacteria cultured from the microbiome associated with maize. PeerJ 9:e11359. doi: 10.7717/peerj.11359.

46. Lee C-T, Chen I-T, Yang Y-T, Ko T-P, Huang Y-T, Huang J-Y, Huang M-F, Lin S-J, Chen C-Y, Lin S-S, Lightner DV, Wang H-C, Wang AH-J, Wang H-C, Hor L-I, Lo C-F (2015) The opportunistic marine pathogen *Vibrio parahaemolyticus* becomes virulent by acquiring a plasmid that expresses a deadly toxin. PNAS 112:10798–10803. doi:10.1073/pnas.1503129112.

47. Letchumanan V, Chan K-G, Lee L-H (2014) *Vibrio parahaemolyticus*: a review on the pathogenesis, prevalence, and advance molecular identification techniques. Front Microbiol 5:705. 10.3389/fmicb.2014.00705.

48. Letchumanan V, Chan K-G, Lee L-H (2015) An insight of traditional plasmid curing in *Vibrio* species. Front Microbiol 6:735. 10.3389/fmicb.2015.00735.

49. Letunic I, Bork P (2021) Interactive Tree of Life (iTOL) v5: an online tool for phylogenetic tree display and annotation. Nucleic Acids Res 49:W293–W296. 10.1093/nar/gkab301.

50. Li P, Xin W, Xia S, Luo Y, Chen Z, Jin D, Gao S, Yang H, Ji B, Wang H, Yan Y, Kang L, Wang J (2018) MALDI-TOF mass spectrometry-based serotyping of *V. parahaemolyticus* isolated from the Zhejiang province of China. BMC Microbiol 18:185. 10.1186/s12866-018-1328-z.

51. Liu D, Zhou L, Zhang Z, Zhang Y, Wang Z, Li S, Zhu Y, Zheng H, Zhang Z, Tian Z (2024) Epidemiological and genomic analysis of *Vibrio parahaemolyticus* isolated from imported travelers at the port of Shanghai, China (2017-2019). BMC Microbiol 24:145. 10.1186/s12866-024-03303-7.

52. Liu T, Kang L, Xu J, Wang J, Gao S, Li Y, Li J, Yuan Y, Yuan B, Wang J, Zhao B, Xin W (2022) PVBase: a MALDI-TOF MS database for fast identification and characterization of potentially pathogenic *Vibrio* species from multiple regions of China. Front Microbiol 13:872825. 10.3389/fmicb.2022.872825.

53. Makino K, Oshima K, Kurokawa K, Yokoyama K, Uda T, Tagomori K, Iijima Y, Najima M, Nakano M, Yamashita A, Kubota Y, Kimura S, Yasunaga T, Honda T, Shinagawa H, Hattori M, Iida T (2003) Genome sequence of *Vibrio parahaemolyticus*: a pathogenic mechanism distinct from that of *V. cholerae*. Lancet 361:743–749. 10.1016/S0140-6736(03)12659-1.

54. Mazhari F, Regberg AB, Castro CL, LaMontagne MG (2025) Resolution of MALDI-TOF compared to whole genome sequencing for identification of *Bacillus* species isolated from cleanrooms at NASA Johnson Space Center. Front Microbiol 16:1499516. 10.3389/fmicb.2025.1499516.

55. McLaughlin JB, DePaola A, Bopp CA, Martinek KA, Napolilli NP, Allison CG, Murray SL, Thompson EC, Bird MM, Middaugh JP (2005) Outbreak of *Vibrio parahaemolyticus* gastroenteritis associated with Alaskan oysters. N Engl J Med 353:1463–1470. 10.1056/NEJMoa051594.

56. Mellmann A, Bimet F, Bizet C, Borovskaya AD, Drake RR, Eigner U, Fahr AM, He Y, Ilina EN, Kostrzewa M, Maier T, Mancinelli L, Moussaoui W, Prévost G, Putignani L, Seachord CL, Tang YW, Harmsen D (2009) High interlaboratory reproducibility of matrix-assisted laser desorption ionization-time of flight mass spectrometry-based species identification of nonfermenting bacteria. J Clin Microbiol 47:3732–3734. doi:10.1128/jcm.00921-09.

57. Miller JJ, Weimer BC, Timme R, Lüdeke CHM, Pettengill JB, Bandoy DJD, Weis AM, Kaufman J, Huang BC, Payne J, Strain E, Jones JL (2021) Phylogenetic and biogeographic patterns of *Vibrio parahaemolyticus* strains from North America inferred from whole-genome sequence data. Appl Envir Microbiol 87:e01403–20. doi:10.1128/AEM.01403-20.

58. Mougin J, Flahaut C, Roquigny R, Bonnin-Jusserand M, Grard T, Le Bris C (2020) Rapid identification of *Vibrio* species of the harveyi clade using MALDI-TOF MS profiling with main spectral profile database implemented with an in-house database: Luvibase. Front Microbiol 11:586536. 10.3389/fmicb.2020.586536.

59. Moussa Pouly M, Cauvin E, Le Piouffle A, Bidault A, Paillard C, Benoit F, Thuillier B, Treilles M, Travers M-A, Garcia C, MALDI-TOF database for the identification of potentially pathogenic Vibrio in marine molluscs, SEANOE.

60. Naka H, Dias GM, Thompson CC, Dubay C, Thompson FL, Crosa JH (2011) Complete genome sequence of the marine fish pathogen *Vibrio anguillarum* harboring the pJM1 virulence plasmid and genomic comparison with other virulent strains of *V. anguillarum* and *V. ordalii*. Infect Immun 79:2889–2900. doi:10.1128/iai.05138-11.

61. Ndraha N, Hsiao H-I (2022) A climate-driven model for predicting the level of *Vibrio parahaemolyticus* in oysters harvested from Taiwanese farms using elastic net regularized regression. Microb Risk Anal 21:100201. 10.1016/j.mran.2022.100201.

62. Nguyen L-T, Schmidt HA, von Haeseler A, Minh BQ (2014) IQ-TREE: A fast and effective stochastic algorithm for estimating maximum-likelihood phylogenies. Mol Biol Evol 32:268–274. 10.1093/molbev/msu300.

63. Page AJ, Cummins CA, Hunt M, Wong VK, Reuter S, Holden MTG, Fookes M, Falush D, Keane JA, Parkhill J (2015) Roary: rapid large-scale prokaryote pan genome analysis. Bioinform 31:3691–3693. 10.1093/bioinformatics/btv421.

64. Parks DH, Imelfort M, Skennerton CT, Hugenholtz P, Tyson GW (2015) CheckM: assessing the quality of microbial genomes recovered from isolates, single cells, and metagenomes. Genome Res 25:1043–1055. 10.1101/gr.186072.114.

65. Pavlovic M, Konrad R, Iwobi AN, Sing A, Busch U, Huber I (2012) A dual approach employing MALDI-TOF MS and real-time PCR for fast species identification within the *Enterobacter cloacae* complex. FEMS Microbiol Lett 328:46–53. 10.1111/j.1574-6968.2011.02479.x.

66. Price V, Ngwira LG, Lewis JM, Baker KS, Peacock SJ, Jauneikaite E, Feasey N (2023) A systematic review of economic evaluations of whole-genome sequencing for the surveillance of bacterial pathogens. Microb Genom 9:000947. 10.1099/mgen.0.000947.

67. Prithvisagar KS, Krishna Kumar B, Kodama T, Rai P, Iida T, Karunasagar I, Karunasagar I (2021) Whole genome analysis unveils genetic diversity and potential virulence determinants in *Vibrio parahaemolyticus* associated with disease outbreak among cultured *Litopenaeus vannamei* (Pacific white shrimp) in India. Virulence 12:1936–1949. 10.1080/21505594.2021.1947448.

68. Rahmani A, Vercauteren M, Vranckx K, Boyen F, Bidault A, Pichereau V, Decostere A, Paillard C, Chiers K (2021) MALDI-TOF MS as a promising tool to assess potential virulence of *Vibrio tapetis* isolates. Aquaculture 530:735729. 10.1016/j.aquaculture.2020.735729.

69. Romeo T, Vakulskas CA, Babitzke P (2013) Post-transcriptional regulation on a global scale: form and function of Csr/Rsm systems. Environ Microbiol 15:313–324. 10.1111/j.1462-2920.2012.02794.x.

70. Rudrik JT, Soehnlen MK, Perry MJ, Sullivan MM, Reiter-Kintz W, Lee PA, Pettit D, Tran A, Swaney E (2017) Safety and accuracy of matrix-assisted laser desorption ionization–time of flight mass spectrometry for identification of highly pathogenic organisms. J Clin Microbiol 55:3513–3529. doi:10.1128/jcm.01023-17.

71. Sawabe T, Ogura Y, Matsumura Y, Gao F, Amin AR, Mino S, Nakagawa S, Sawabe T, Kumar R, Fukui Y, Satomi M, Matsushima R, Thompson FL, Gomez-Gil B, Christen R, Maruyama F, Kurokawa K, Hayashi T (2013) Updating the *Vibrio* clades defined by multilocus sequence phylogeny: proposal of eight new clades, and the description of *Vibrio tritonius* sp. nov. Front Microbiol 4:00414. 10.3389/fmicb.2013.00414.

72. Schwengers O, Jelonek L, Dieckmann MA, Beyvers S, Blom J, Goesmann A (2021) Bakta: rapid and standardized annotation of bacterial genomes via alignment-free sequence identification. Microb Genom 7:000685. 10.1099/mgen.0.000685.

73. Seemann T (2014) Prokka: rapid prokaryotic genome annotation. Bioinform 30:2068–2069. 10.1093/bioinformatics/btu153.

74. Sekiguchi Y, Teramoto K, Tourlousse DM, Ohashi A, Hamajima M, Miura D, Yamada Y, Iwamoto S, Tanaka K (2023) A large-scale genomically predicted protein mass database enables rapid and broad-spectrum identification of bacterial and archaeal isolates by mass spectrometry. Genome Biol 24:257. 10.1186/s13059-023-03096-4.

75. Semenza JC, Trinanes J, Lohr W, Sudre B, Löfdahl M, Martinez-Urtaza J, Nichols GL, Rocklöv J (2017) Environmental suitability of *Vibrio* infections in a warming climate: An early warning system. Environ Health Perspect 125:107004. 10.1289/ehp2198.

76. Shimodaira H (2002) An approximately unbiased test of phylogenetic tree selection. Syst Biol 51:492–508. 10.1080/10635150290069913.

77. Sibińska E, Arendowski A, Fijałkowski P, Gabryś D, Pomastowski P (2024) Comparison of the Bruker Microflex LT and Zybio EXS2600 MALDI TOF MS systems for the identification of clinical microorganisms. Diagn Microbiol Infect Dis 108:116150. 10.1016/j.diagmicrobio.2023.116150.

78. Singhal N, Kumar M, Kanaujia PK, Virdi JS (2015) MALDI-TOF mass spectrometry: an emerging technology for microbial identification and diagnosis. Front Microbiol 6:e791. doi: 10.3389/fmicb.2015.00791.

79. Souvorov A, Agarwala R, Lipman DJ (2018) SKESA: strategic k-mer extension for scrupulous assemblies. Genom Biol 19:153. 10.1186/s13059-018-1540-z.

80. Stephan R, Ziegler D, Pflüger V, Vogel G, Lehner A (2010) Rapid genus- and species-specific identification of *Cronobacter* spp. by matrix-assisted laser desorption ionization-time of flight mass spectrometry. J Clin Microbiol 48:2846–2851. 10.1128/jcm.00156-10.

81. Suarez S, Ferroni A, Lotz A, Jolley KA, Guérin P, Leto J, Dauphin B, Jamet A, Maiden MCJ, Nassif X, Armengaud J (2013) Ribosomal proteins as biomarkers for bacterial identification by mass spectrometry in the clinical microbiology laboratory. J Microbiol Method 94:390–396. 10.1016/j.mimet.2013.07.021.

82. Sulaiman IM, Miranda N, Hook W, Mendoza J, Kumfert Q, Barnes T, Sung K, Khan S, Nawaz M, Banerjee P, Simpson S, Karem K (2023) A single-laboratory performance evaluation of MALDI-TOF MS in rapid identification of *Staphylococcus aureus*, *Cronobacter sakazakii*, *Vibrio parahaemolyticus*, and some closely related bacterial species of public health importance. J AOAC Int 106:1574–1588. 10.1093/jaoacint/qsad109.

83. Suzuki R, Shimodaira H (2006) Pvclust: an R package for assessing the uncertainty in hierarchical clustering. Bioinformatics 22:1540–1542. doi: 10.1093/bioinformatics/btl117.

84. Tan KE, Ellis BC, Lee R, Stamper PD, Zhang SX, Carroll KC (2012) Prospective evaluation of a matrix-assisted laser desorption ionization–time of flight mass spectrometry system in a hospital clinical microbiology laboratory for identification of bacteria and yeasts: a bench-by-bench study for assessing the impact on time to identification and cost-effectiveness. J Clin Microbiol 50:3301–3308. doi:10.1128/jcm.01405-12.

85. Tempel S, Using and Understanding RepeatMasker. in: Y. Bigot, (Ed.), Mobile Genetic Elements: Protocols and Genomic Applications, Humana Press, Totowa, NJ, 2012, pp. 29–51.

86. Timme RE, Pfefer T, Bias CH, Allard MW, Huang X, Strain E, Balkey M, A comprehensive guide to quality assessment and data submission for genomic surveillance of enteric pathogens. in: A. Bridier, (Ed.), Foodborne Bacterial Pathogens: Methods and Protocols, Springer US, New York, NY, 2025, pp. 199–209.

87. Topić Popović N, Kazazić SP, Bojanić K, Strunjak-Perović I, Čož-Rakovac R (2023) Sample preparation and culture condition effects on MALDI-TOF MS identification of bacteria: A review. Mass Spectrom Rev 42:1589–1603. 10.1002/mas.21739.

88. Trinanes J, Martinez-Urtaza J (2021) Future scenarios of risk of *Vibrio* infections in a warming planet: a global mapping study. Lancet Planet Health 5:e426–e435. 10.1016/S2542-5196(21)00169-8.

89. Vargha M, Takáts Z, Konopka A, Nakatsu CH (2006) Optimization of MALDI-TOF MS for strain level differentiation of *Arthrobacter* isolates. J Microbiol Method 66:399–409. 10.1016/j.mimet.2006.01.006.

90. Vezzulli L, Grande C, Reid PC, Hélaouët P, Edwards M, Höfle MG, Brettar I, Colwell RR, Pruzzo C (2016) Climate influence on *Vibrio* and associated human diseases during the past half-century in the coastal North Atlantic. PNAS 113:E5062–E5071. doi:10.1073/pnas.1609157113.

91. Vidal LMR, Venas TM, Gonçalves ARP, Mattsson HK, Silva RVP, Nóbrega MS, Azevedo GPR, Garcia GD, Tschoeke DA, Vieira VV, Thompson FL, Thompson CC (2020) Rapid screening of marine bacterial symbionts using MALDI-TOF MS. Arch Microbiol 202:2329–2336. 10.1007/s00203-020-01917-9.

92. Wick RR, Judd LM, Gorrie CL, Holt KE (2017) Unicycler: resolving bacterial genome assemblies from short and long sequencing reads. PLoS Comput Biol 13:e1005595. doi: 10.1371/journal.pcbi.1005595.

