## Supplementary figures and images for "Resolution of MALDI-TOF MS Compared to Whole Genome Sequencing for the Identification of *Vibrio parahaemolyticus* Strains Isolated from Oysters"

### S4.tiff

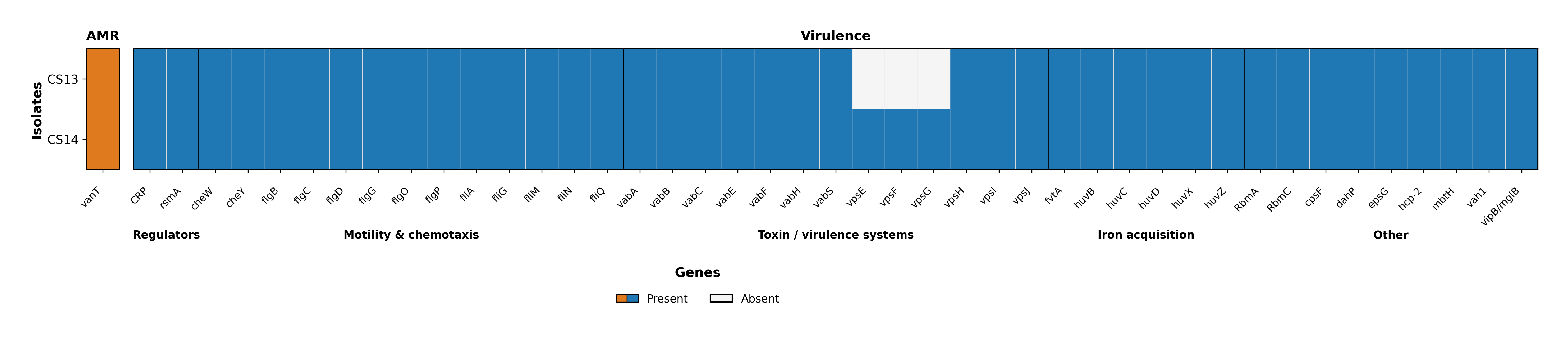
